# Identification of the *Col4a3*^-/-^ mouse as a model of chronic kidney disease-associated cachexia

**DOI:** 10.64898/2026.10.05.756697

**Authors:** Marie Buléon, Manon Brunet, Emma Ballongue, Victoire Ankaoua, Elsa Gutowski, Laure Monteillet, Rayane Benyahia, Guylène Feuillet, Stéphane Poitevin, Stéphane Burtey, Laetitia Koppe, Dominique Langin, Maria Rohm, Joost P. Schanstra, Julie Klein, Pauline Morigny

## Abstract

**Background:** Cachexia is a frequent complication of advanced chronic kidney disease (CKD), *i.e.* renal cachexia, particularly in patients undergoing hemodialysis in whom it represents one of the strongest predictors of morbidity and mortality. Yet, our understanding of the molecular mechanisms leading to body wasting in CKD and management of cachectic patients are still poorly defined. To date, a preclinical mouse model of robust and reproducible CKD-associated cachexia is missing. Here, we assessed in a systematic manner the cachectic phenotype of the genetic *Col4a3^-/-^* mouse model of CKD.

**Methods:** 129/SvJ *Col4a3^-/-^* male and female animals were monitored longitudinally until they developed kidney failure. Measured glomerular filtration rate (mGFR), body composition and grip strength were recorded at baseline (6 weeks of age, before onset of kidney dysfunction) and at endpoint (CKD with weight loss or 12 weeks of age). The severity of CKD was confirmed by biochemical and histological analyses. Muscle atrophy was assessed via fiber cross-sectional area analysis, atrogene expression and transcriptomics. Adipose tissue wasting was assessed by measuring adipocyte area and proteins involved in lipid synthesis and breakdown. Liver acute phase response and systemic inflammation were evaluated by measuring circulating levels of key factors using ELISA.

**Results:** *Col4a3^-/-^* animals developed CKD (decrease in mGFR: -85%, P<0.0001) and cachexia within 8-12 weeks of age. Cachexia was characterized by a progressive loss of body weight (-17.8% ±1.0, P<0.0001), lean (-8.2% ±2.0, P<0.0001) and fat mass (-66.5% ±4.7, P<0.0001) associated with reduced grip strength (-24.3% ±0.0, P=0.0002). At the tissue level, cachexia led to muscle atrophy and induction of atrogenes. Muscle transcriptomics further validated the cachectic phenotype of the *Col4a3^-/-^*, with 47% of genes altered in a similar manner with the well-established colon-C26 mouse model of cancer cachexia. It also revealed signatures that appeared more specific to renal cachexia. Adipose tissue wasting was characterized by a depletion in fat stores, consecutive to an imbalance between proteins involved in lipogenesis and lipolysis. Cachexia in this model was also associated with the induction of acute phase response in liver, elevated circulating levels of serum amyloid A and systemic inflammation, with the induction of established cachexia mediators (IL-6, GDF-15 P<0.0001). Both male and female mice developed cachexia.

**Conclusions:** Our data highlight the potential of using the *Col4a3^-/-^*mouse model of CKD for the future study of renal cachexia, offering a more robust alternative to classically used models of partial nephrectomy or adenine-rich diet.

## Introduction

Cachexia is a devastating wasting syndrome, characterized by an involuntary loss of body weight, lean and/or fat mass, that cannot be solely reversed by nutritional supplementation^1^. Cachectic patients experience frailty, have a poor quality of life and tolerance to therapies, leading to increased hospitalization and mortality rates. It often affects patients at advanced stages of chronic diseases, such as cancer, chronic kidney, heart and pulmonary diseases. General cachexia features include activation of proteolytic pathways in skeletal muscle leading to atrophy (*e.g.* activation of the ubiquitin-proteasome system, induction of autophagy markers), pro-lipolytic pathways in adipose tissue, acute phase response in the liver, anorexia and a sustained inflammatory state. While most clinical and experimental studies to date have focused on cancer-associated cachexia, other forms of cachexia remain comparatively understudied. This is particularly true for chronic kidney disease (CKD)-associated cachexia (*i.e.* renal cachexia), despite its high prevalence and major contribution to morbidity and mortality in patients with advanced CKD^2–4^.

CKD is a serious problem of public health, affecting > 850 million individuals worldwide and representing the 9^th^ leading cause of premature death with a considerable increase in the past 30 years (World Health Organization)^5^. CKD is defined by the presence of kidney damage or an estimated glomerular filtration rate (GFR) of less than 60 mL/min/1.73m², persisting for a minimum of 3 months. Hallmarks of CKD include blood (accumulation of metabolic waste in the blood *e.g.* urea) and urine abnormalities (proteinuria or hematuria) and histological lesions (*e.g.* glomerulosclerosis, tubular injury, interstitial fibrosis). Patients with end-stage CKD (*i.e.* kidney failure) and kidney replacement therapy (*i.e.* dialysis or transplantation) are particularly at risk for malnutrition and tissue wasting, referred as protein energy wasting syndrome or kidney disease/renal cachexia at the most advanced stages^2,4,6^. Different definitions of renal cachexia have been proposed over time^2,7–9^. Nevertheless, all rely on phenotypic features, including unintentional weight loss > 5 % over a period of 6-12 months, low body mass index, reduced skeletal muscle mass and/or fat mass, together with etiologic criteria such as reduced dietary intake. Renal cachexia is one of the strongest predictors of morbidity, mortality and emotional distress in patients with CKD^2–4^. Treatment options for renal cachexia are yet poorly defined and prove mostly inefficient^1,4,6,10^.

Limitations for the study of renal cachexia reside in part in the available animal models of CKD. Historically, the model of 5/6 nephrectomy, in which one kidney and two-thirds of the contralateral kidney are surgically removed^11^, and the model of adenine-rich diet, which promote crystal deposition within kidney tubules^12^, are the most widely-used for the study of renal cachexia. Those models have been particularly useful for identifying primary CKD-related factors that drive metabolic alterations in skeletal muscle, adipose tissue or liver^10,13–17^. However, mechanistic studies of renal cachexia may be mitigated by the relatively mild cachectic phenotype observed in these animals. In addition, these models are experimentally-induced and may be affected by confounding factors, including surgical stress or adenine overload, both of which can independently influence energy metabolism. Additional models of CKD displaying robust and reproducible cachectic phenotypes are therefore warranted to advance mechanistic research in renal cachexia.

Beyond these models, several other models of CKD have been developed, including genetically-driven models that reproduce specific causes or trajectories of CKD. However, in many of these models, cachexia has not been systematically assessed. As a result, their relevance for the study of renal cachexia remains largely underexplored. Among these models, the well-established *Col4a3*^-/-^ mouse model of Alport syndrome represents a particularly relevant genetic model of progressive CKD^18^. In patients with Alport syndrome, mutations in *COL4A3*, *COL4A4*, or *COL4A5* disrupt the type IV collagen network of the glomerular basement membrane, leading to progressive CKD development and, in severe forms (*e.g.* X-linked), kidney failure requiring kidney replacement therapy at a young age, with a reported mean age of 22 years^19^. Despite its broad use to study CKD progression and related complications, the cachectic phenotype of this model has not been fully characterized.

Here, we have performed a systematic characterization of *Col4a3*^-/-^ mouse model in regards to cachexia development. In our study, we performed a longitudinal follow-up of body composition, muscle strength, and systematic assessment of established cachexia markers in male and female *Col4a3*^-/-^ mice. Our data showed that *Col4a3*^-/-^ animals develop cachectic phenotypes with features similar to those observed in cancer cachexia models, highlighting the potential of this model for mechanistic studies of renal cachexia.

## Methods

### Animal experiments

Female and male *Col4a3^+/-^* heterozygous mice on 129/SvJ genetic background were obtained from The Jackson Laboratory (129-*Col4a3^tm1Dec^*/J, strain #002908, RRID:IMSR_JAX:002908) and bred at the *Centre Régional d’Exploration Fonctionnelle et Ressources Expérimentales* (Toulouse, France). Descendants were genotyped to select heterozygous and *Col4a3^-/-^* homozygous knock-out mice (primers used for genotyping: gen-*Col4a3* For: 5’-ATAGCCCACTCGTCTCTAAGAT-3’; gen-*Col4a3* Rev : 5’-GGAAGCTCGAATCCTATAAATAGTGTA-3’; gen-NEO For : 5’-GATACCGTAAAGCACGAGGAAG-3’; gen-NEO Rev : 5’-TCAGGATGATCTGGACGAAGA-3’). Animals were housed with unlimited access to water and food (mouse standard chow diet Safe Diets #A04), and kept on a 12-hour light–dark cycle, at 22 °C, in a pathogen-free environment.

Mouse condition was assessed daily. Body weight was recorded bi-weekly while animals were stable and then daily once they started to lose weight. Animals were necropsied once they reached experimental endpoint (body weight loss > 15 %, bad general condition or age of 12 weeks). Measured GFR (mGFR), body composition and grip strength were measured at 6 weeks of age (initial measurements) and at endpoint. Mice were euthanized by an overdose of pentobarbital after isoflurane sedation. Blood was withdrawn from the vena cava, transferred into heparin tubes (Microvette, Sarsdedt # 20.1345) and centrifuged at 2,500 *x g*, 4°C for 5 min. Tissues were immediately collected, snap frozen, or fixed in PFA for further analysis.

### Glomerular filtration rate

mGFR was measured by using a transdermal fluorescence detector (MediBeacon) placed on the epilated back skin of mice anesthetized with isoflurane. Baseline auto-fluorescence was determined during the first 1-minute measurement. The animals received sinistrin-FITC (7 mg/100 g of body weight, Medibeacon INC #FTCFS001) by retro-orbital injection and placed alone in a cage without access to water, and fluorescence was measured for 45 minutes. Fluorescence data were analyzed with the MPD Lab software (v2.15, Medibeacon). Data were normalized to the mouse body weight.

### Body composition

Body composition was measured by nuclear magnetic resonance with a Bruker’s minispec Whole Body Composition Analyzer.

### Grip strength

Grip strength was recorded at the beginning and at the end of experiment. Mice were held by the root of the tail and lowered over a grid connected to a force sensor (Bioseb Instruments # BIO-GS3), where they immediately held on with the front paws. As the animals were gently pulled away from the grid in a horizontal manner, the maximal force with which mice could hold on was recorded. Such measurement was carried out three times in succession, with resting times between two recordings, and we used the mean of all values.

### Histological staining

Kidneys, gastrocnemius muscles and subcutaneous adipose tissues were fixed in paraformaldehyde 4% and embedded in paraffin. 4 µm-thick sections were cut and mounted on glass slides. Slides were deparaffinized and stained with either Masson’s Trichrome (Goldner Trichrome, Diapath #010224), Sirius Red (Sirius Red 0.1% in saturated picric acid, Electron Microscopy Science # 2635702) or Hemalun/Eosin (Dako Mayer’s Hematoxylin (Lillie’s Modification) #S3309); Eosin 2%) according to manufacturers’ instructions. Slides were then digitalized with a Nanozoomer 2.0 RS (Hamamatsu Photonics SARL).

Fibrosis was quantified on kidney and muscle sections stained with Sirius Red using Image J software and expressed as % of total section area.

For kidneys, tubular injury was blindly scored on Masson trichrome-stained sections by 3 investigators following the criteria: 0: no tubular injury; 1: <10% of injured tubules; 2: 10-24% of injured tubules; 3: 25-49% of injured tubules; 4: 50-74% of injured tubules; 5: >75% of injured tubules. The score considers tubular dilatation, tubular atrophy, tubular cast formation, sloughing of tubular epithelial cells, loss of the brush border, or thickening of the tubular basement membrane. The percentage of sclerotic glomeruli was determined by randomly selecting 20 to 40 glomeruli per section and counting the number of sclerotic glomeruli among them.

For muscle and adipose tissue, quantification of myofibers and adipocyte cross-sectional area was performed using QuPath, with automatic fiber/cell selection (using a size threshold between 400 and 2500 µm^2^ and circularity threshold at 0.42 for muscle fibers; and size threshold between 300 and 2500 µm^2^ for adipocytes). ≥ 230 muscle fibers and ≥ 150 adipocytes per animal were quantified.

### Plasma analyses

Blood urea was measured on a Pentra 400 analyzer (Horiba Medical) by the Phenotyping Department of the Anexplo platform (CREFRE, Toulouse, France). Plasma levels of serum amyloid A (SAA), interleukin-6 (IL-6) and growth differentiation factor-15 (GDF-15) were determined using ELISA kits (R&D Systems #MSAA00, #M6000B, #MGD150) according to manufacturer’s instructions. Dilution of plasma samples for detection was optimized to 1/1,000 (SAA), 1/5 (IL-6) and 1/10 (GDF-15).

### Urinary analyses

Urine samples were collected at endpoint. Hematuria was measured using test strips (#Multistix 8SG SIEMENS). Proteinuria was analyzed using the Pentra 400 (HORIBA Medical) at the Phenotyping Department of the Anexplo platform (CREFRE, Toulouse, France).

### qPCR analysis

Pieces of tissues were lysed in Trizol (Life Technologies #15596018). RNA was then isolated using chloroform, precipitated with isopropanol and washed in 75 % ethanol. RNA concentration was assessed using the #Nanodrop 2000 (Thermo Fisher Scientific). RNA was treated with DNase I (Thermo Fisher Scientific #18068-015) before performing cDNA synthesis (Life technologies #4368814). qPCR analysis was perfomed using Thermo Fisher QuantStudio 5 Real-Time PCR system and Owi Green FAST qPCR Premix (Owibio #OZYA008). Specific primers were obtained from Origene (*Foxo3* #MP204949, *Fbxo32* #MP204713, *Trim63* #MP217600, *Gabarapl1* #MP205204, *Map1lc3b* #MP208170, *Ctsl* #MP201834, *Bnip3* #MP201275, *Lcn2* # MP207571, *Fn1* #MP204834, *Crp* #MP202974, *Tbp* #MP216640, *Hprt* #MP206455) except from *Fga* (F: GGATTCTAACTCACTGACCAGGA; R: CCTCAGGATCTCAATTCTGCGC) and *Saa1/2* (F: GGAGTCTGGGCTGCTGAGAAAA; R: TGTCTGTTGGCTTCCTGGTCAG). Gene expression was normalized to *Tbp* or *Hprt* mRNA levels. 2^-ΔCt^ values were used for statistical analysis.

### Western blot

Tissues were lysed in ice-cold lysis buffer (Tris pH 8 2 mM, EDTA 5 mM, NaCl 150 mM, NP-40 0.5 %) with 1 X cOmplete protease inhibitor cocktail and 1 X PhosSTOP cocktail (Sigma-Aldrich (Roche) # 11836170001, # 4906837001). Proteins were loaded onto tris-glycine gels (Bio-rad #4568086) and transferred onto nitrocellulose membranes (Cytiva #10600002) using the Trans-Blot Turbo Transfer System (Bio-rad). After blocking, membranes were incubated with the following primary antibodies: ACC (Cell Signaling Technology, CST #3662, 1/1,000), ATGL (CST #2138, 1/1,000) and HSL (CST #4107, 1/1,000). After incubation with anti-rabbit secondary antibody coupled to horse radish peroxidase (CST #7074, 1/10,000), protein levels were determined using the ECL western blotting reagents (Cytivia #RPN2209) and the ChemiDoc MP System (Bio-rad). Signals were quantified using ImageLab software (Bio-rad) and normalized to Ponceau staining (Sigma-Aldrich #P7170).

### RNA-sequencing

RNA-sequencing of tibialis anterior was performed at Novogene (Munich, Germany). RNA was extracted using columns (Econospin #1920-250).

*Library Construction, Quality Control and Sequencing:* Messenger RNA was purified from total RNA using poly-T oligo-attached magnetic beads. After fragmentation, the first strand cDNA was synthesized using random hexamer primers, followed by the second strand cDNA synthesis using either dUTP for directional library or dTTP for non-directional library. Non-directional library was ready after end repair, A-tailing, adapter ligation, size selection, amplification, and purification. Directional library was ready after end repair, A-tailing, adapter ligation, size selection, USER enzyme digestion, amplification, and purification. The library was checked with Qubit and real-time PCR for quantification and bioanalyzer for size distribution detection. Quantified libraries were pooled and sequenced on Illumina platforms, according to effective library concentration and data amount.

*Clustering and sequencing:* After library quality control, different libraries were pooled based on the effective concentration and targeted data amount, then subjected to Illumina sequencing.

*Bioinformatics Analysis Pipeline:*

*1) Data quality control:* Raw data (raw reads) of fastq format were firstly processed through fastp software. In this step, clean data (clean reads) were obtained by removing reads containing adapter, reads containing ploy-N and low-quality reads from raw data. At the same time, Q20, Q30 and GC content the clean data were calculated. All the downstream analyses were based on the clean data with high quality.
*2) Reads mapping to the reference genome:* Reference genome (mm39) and gene model annotation files were downloaded from genome website. Use HISAT2 (2.2.1) to build the index of the reference genome, and use HISAT2 to align paired-end clean reads to 2 the reference genome. HISAT2 can use the gene model annotation file to create splice-aware alignments, providing better alignment accuracy compared to other non-splice alignment tools.
*3) Quantification of gene expression level:* featureCounts (2.0.6) was used to count the reads numbers mapped to each gene. Then FPKM of each gene was calculated based on the length of the gene and reads count mapped to this gene.
*4) Differential expression analysis*: Differential expression analysis for two groups was performed using the DESeq2 R package (1.42.0). The resulting P-value is adjusted using the Benjamini and Hochberg’s methods to control the error discovery rate. The threshold of significant differential expression was padj <= 0.05 and |log2(foldchange)| >= 1. PCA analysis was performed on the gene expression value (FPKM) of all samples.
*5) Ingenuity Pathway Analysis*: Canonical pathway analysis and upstream regulator analysis were performed using the differentially expressed genes (DEGs) provided by Novogene (padj <= 0.05 and |log2(foldchange)| >= 1) and the Ingenuity Pathway Analysis software (IPA, Qiagen). Comparative pathway analysis was performed using a publicly available dataset, also generated by Novogene and analyzed in a similar manner, from animals with cancer cachexia^20^ (C26 colon carcinoma model, including all genes with padj <= 0.05). In Figure 7D, we selected the top significant pathways with concordant z-scores between groups.

### Statistical analysis

Except from RNA-sequencing data, statistical analysis was performed using GraphPad Prism 10. Unpaired two-way ANOVA with Sidak’s post-hoc tests were used to compare two variables (group effect x sex effect). Correlation between two variables was assessed using simple linear regression analysis with 95 % confidence interval. Muscle fiber and adipocyte distributions were calculated using the *Frequency distribution* function with a bin width of 100. All quantitative results are presented as means ± standard error of the mean (s.e.m). The exact number of animals studied for each experiment is specified in Figure legends.

## Results

### Glomerular Col4a3 deletion leads to kidney failure and progressive body wasting within 3 months

We longitudinally followed-up *Col4a3*^-/-^ male and female animals until they developed kidney failure (characterized by a decrease in mGFR > 85 % compared to control group) and reached experiment endpoint (body condition, age of 12 weeks) (**Figure 1**). As necropsy was not performed on the same day for all *Col4a3*^-/-^ animals, matching controls were euthanatized at a same age to allow for direct comparison of final tissue weights. Heterozygous *Col4a3^+/-^* animals were used as controls (CTRL) since they were identical in terms of kidney function, kidney weight and body weight compared with *Col4a3^+/+^* (**Figure S1**).

**Figure 1:**
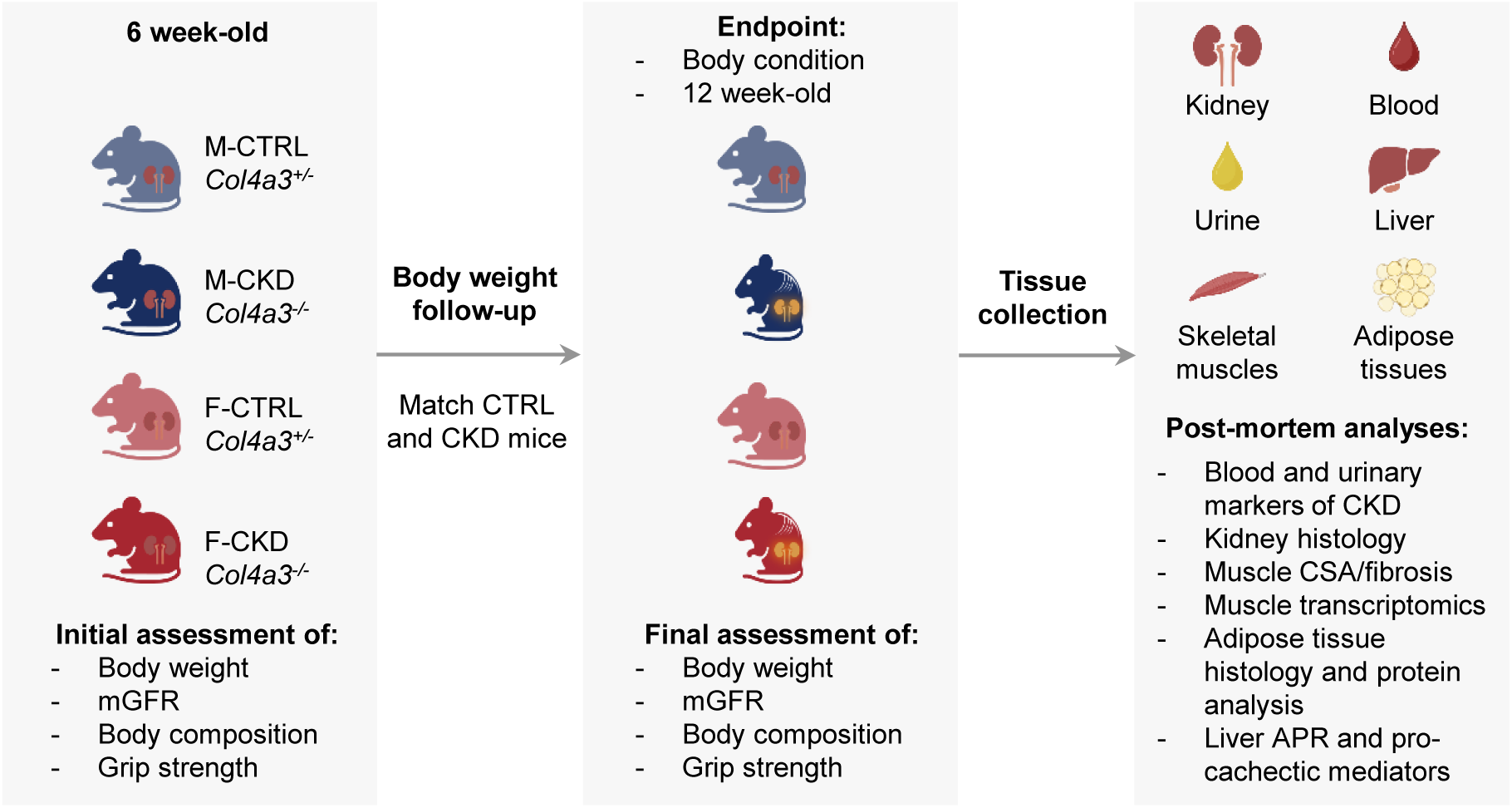
Overview of the experiment. Male (M, blue) and female mice (F, red), heterozygous (CTRL) or knockout for *Col4a3* (CKD) were included in the experiment at the age of 6 weeks and initial body weight, kidney function, body composition and grip strength were recorded. Animals were then monitored until they reached experimental endpoint (deterioration of body condition or age of 12 weeks). After final *in vivo* assessments, tissues relevant for the study of CKD and cachexia were collected for further post-mortem characterization of the animals’ cachectic phenotype. CSA: cross-sectional area. APR: acute phase response.

At baseline, 6-week-old *Col4a3*^-/-^ animals were mostly comparable to CTRL animals, with similar kidney function, body weight and lean mass, a mild reduction in fat mass (-25 %) and identical grip strength (**Figure S2**). *Col4a3*^-/-^ animals developed kidney failure within 3 months, characterized by a strong impairment of mGFR and elevated levels of blood and urinary markers of glomerular dysfunction including blood urea nitrogen, proteinuria and hematuria compared to CTRL (**Figures 2A-C, Figure S3A**). Post-mortem analyses confirmed severe glomerulosclerosis, tubular injury and interstitial fibrosis (**Figures 2D-E, Figure S3B**). Of note, males and females were fully comparable in terms of kidney dysfunction and injury at endpoint.

**Figure 2:**
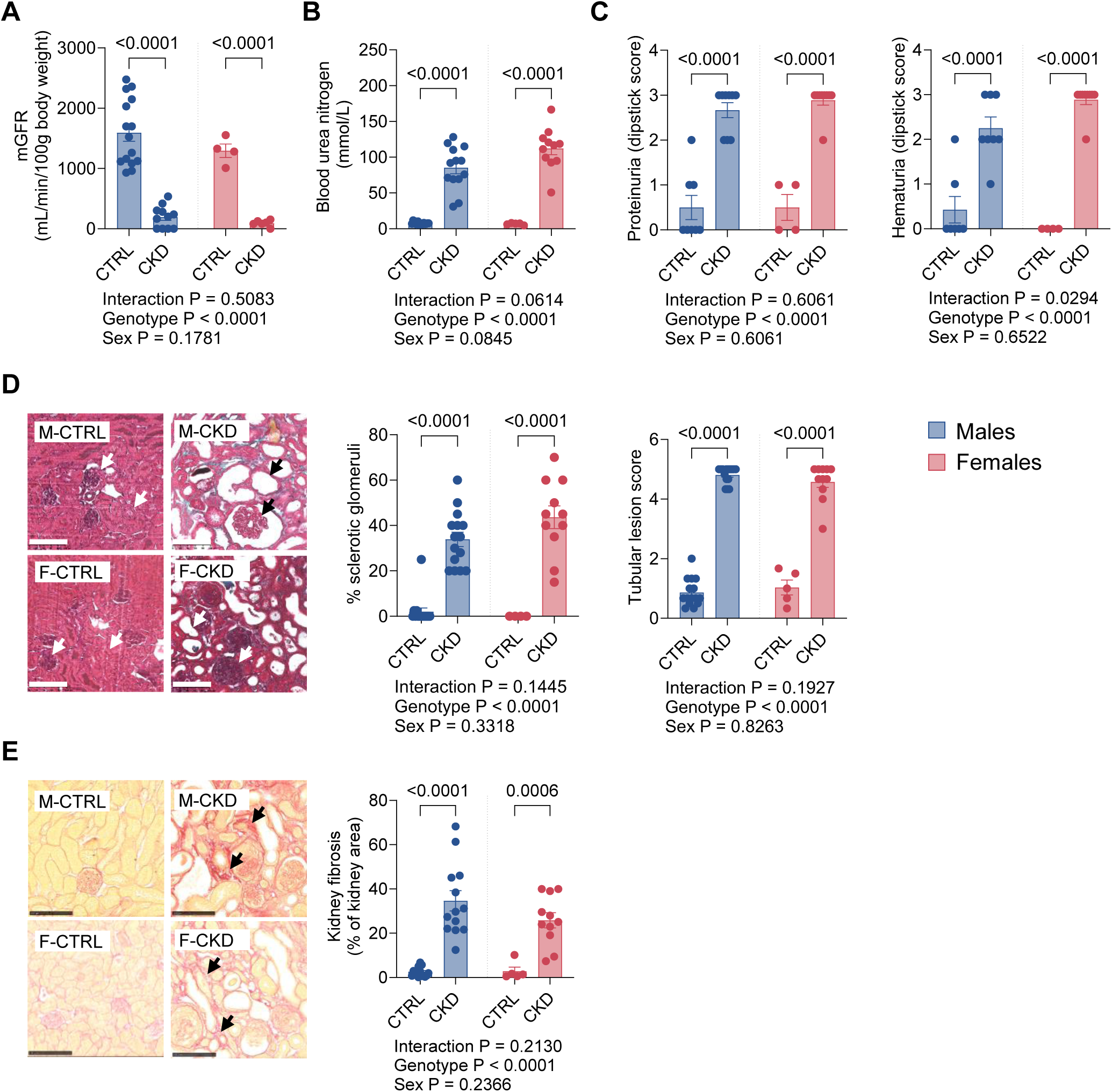
Glomerular *Col4a3* deletion leads to kidney failure within 3 months. Male (M, blue) and female mice (F, red), heterozygous (CTRL) or knockout for *Col4a3* (CKD), were monitored for a maximum of 12 weeks (unless stated otherwise, M-CTRL: n=16, M-CKD: n=15, F-CTRL: n=5, F-CKD: n=11). **(A)** mGFR (M-CTRL: n=15, M-CKD: n=11, F-CTRL: n=4, F-CKD: n=6), **(B)** blood urea nitrogen (M-CTRL: n=15, M-CKD: n=13) and **(C)** urinary markers of CKD (proteinuria, hematuria; M-CTRL: n=7- 8, M-CKD: n=8-9, F-CTRL: n=4, F-CKD: n=9) at endpoint. **(D-E)** Histological assessment (M-CTRL: n=15, M-CKD: n=13-15) of **(D)** sclerotic glomeruli, tubular lesions (arrows show healthy vs damaged glomerules and tubules) and **(E)** kidney fibrosis (Sirius red, arrows show fibrotic lesions). Scale: 100 µm. Data are mean ± s.e.m. Statistical analysis: unpaired two-way ANOVA with Sidak’s post-hoc tests. P values are displayed on the graphs. P < 0.05 was considered significant.

At the same time as kidney failure developed, *Col4a3*^-/-^ animals progressively lost weight, reaching a body weight loss > 15 % within 2-3 weeks (**Figures 3A-B**) with a similar progression between males and females. Body weight loss was inversely correlated with kidney function (**Figure 3C**). To further confirm this relationship between kidney function and body wasting, we performed an additional analysis including only *Col4a3*^-/-^ animals at different stages of CKD, ranging for normal mGFR (around 1000 mL/min/100 g body weight) to kidney failure. We observed again a strong negative correlation between kidney function and the percentage of body weight loss across all knock-out mice, indicating that animals with more severe kidney disease also exhibited greater weight loss (**Figure S4**). Body wasting was associated with the loss of both lean (-8 %) and fat mass (-75 %) compared to CTRL (**Figures 3D-E**).

**Figure 3:**
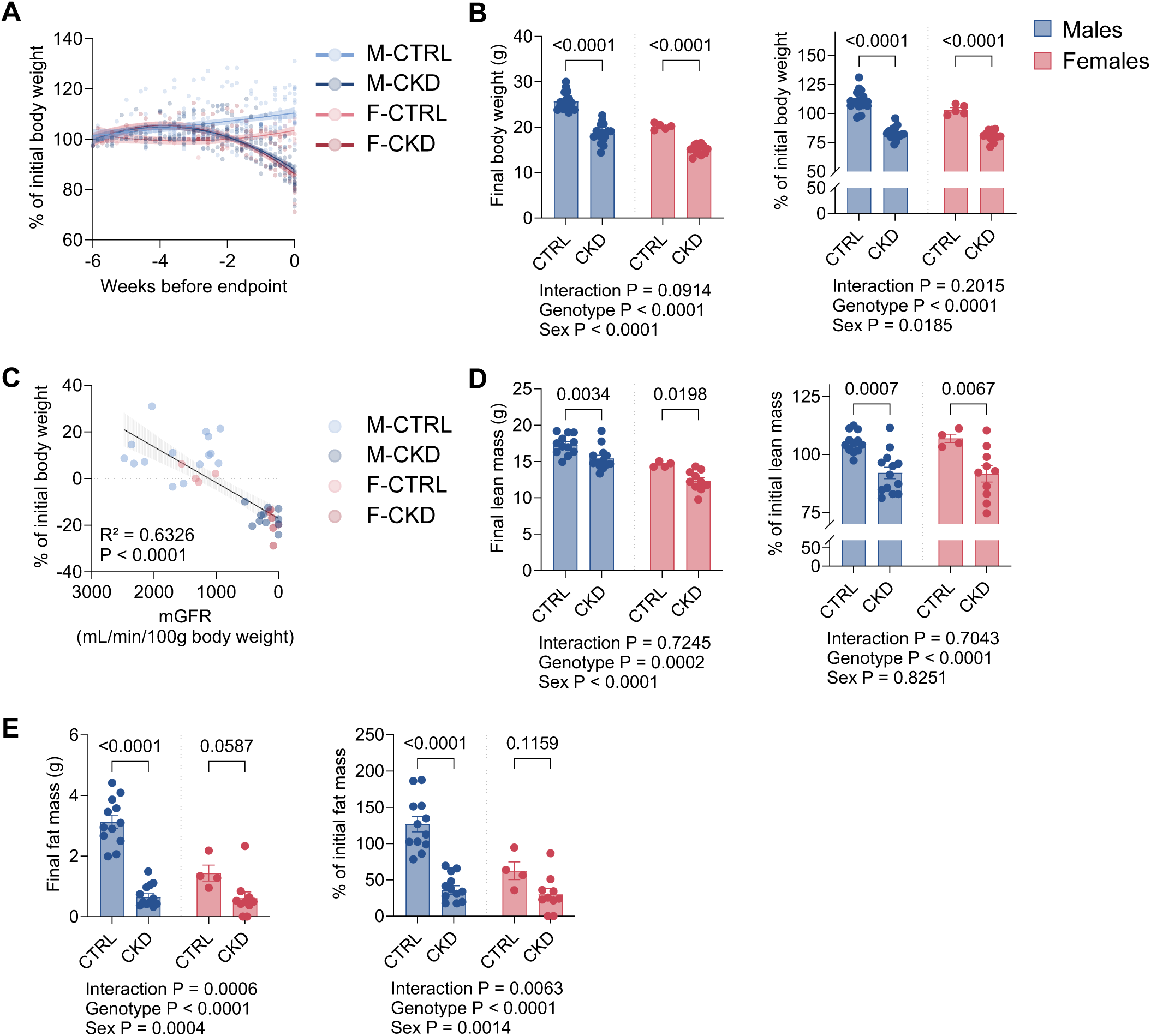
Kidney failure induces body wasting in *Col4a3*-deficient mice. Male (M, blue) and female mice (F, red), heterozygous (CTRL) or knockout for *Col4a3* (CKD), were monitored for a maximum of 12 weeks (unless stated otherwise, M-CTRL: n=16, M-CKD: n=15, F-CTRL: n=5, F-CKD: n=11). **(A)** Kinetic of body weight change throughout the experiment. **(B)** Body weight at endpoint and percentage of body weight loss. **(C)** Association between body weight loss and kidney function (M-CTRL: n=15, M- CKD: n=11, F-CTRL: n=4, F-CKD: n=6). **(D)** Lean mass at endpoint and percentage of lean mass loss (M-CTRL: n=12, M-CKD: n=13, F-CTRL: n=4, F-CKD: n=10). **(E)** Fat mass at endpoint and percentage of fat mass loss (M-CTRL: n=12, M-CKD: n=13, F-CTRL: n=4, F-CKD: n=10). Data are mean ± s.e.m. Statistical analysis: unpaired two-way ANOVA with Sidak’s post-hoc tests (B, D-E), simple linerar regression analysis (C). P values are displayed on the graphs. P < 0.05 was considered significant.

Using publicly available atlas of gene expression (Mouse Cell Atlas^21–23^ and Single Cell Portal^24^), we confirmed that *Col4a3* is expressed in kidney cell populations, but not in metabolic cell types affected by cachexia such as hepatocytes, muscle cells or adipocytes (**Figure S5**). These observations suggest that body wasting in *Col4a3^-/-^*mice is unlikely to result from a direct effect of *Col4a3* deficiency in metabolic tissues, but rather reflects systemic consequences of progressive kidney failure.

Overall, our results indicate that, in this model of genetic nephropathy, cachexia develops in parallel with kidney disease and is not evident prior the onset of kidney failure.

### Kidney failure is associated with muscle atrophy and induction of proteolysis in Col4a3-deficient mice

Muscle wasting is a key determinant of functional impairment in cachectic patients. We therefore assessed classical markers of muscle mass and function in the *Col4a3^-/-^* model of CKD. In this model, kidney failure led to the atrophy of different skeletal muscles compared to CTRL (**Figure 4A**), together with a reduction in muscle grip strength (**Figure 4B**). Muscle atrophy was further confirmed by measuring the cross-sectional area of muscle fibers in gastrocnemius, with a shift towards smaller fibers in animals with kidney failure compared to CTRL (**Figure 4C**). Histologically, muscle atrophy was also accompanied by increased muscle fibrosis (**Figure 4D**). At the molecular level, cachexia was associated with the induction of classical proteolytic markers (*n.b.* ubiquitin ligases and autophagy markers) such as the ubiquitin ligases MurF-1 (*Trim63*) and Atrogin-1 (*Fbxo32*) (**Figures 4E-F**), supporting ongoing muscle proteolysis. Statistical analyses (interaction genotype x sex) did not reveal a major sex effect on muscle wasting. Cachexia in this model was therefore associated with classical markers of muscle atrophy.

**Figure 4:**
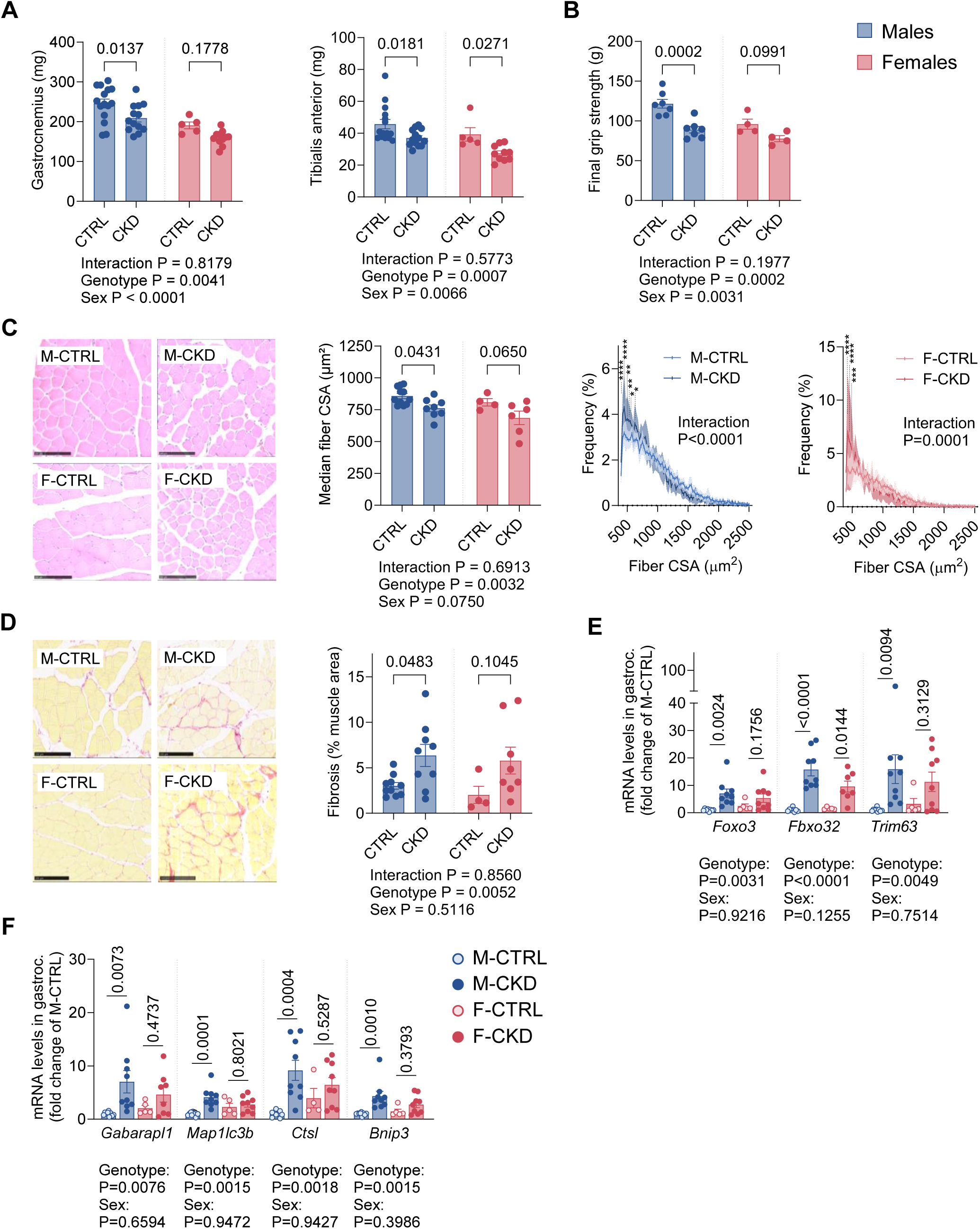
Kidney failure is associated with muscle atrophy and induction of proteolysis in *Col4a3*-deficient mice. Male (M, blue) and female mice (F, red), heterozygous (CTRL) or knockout for *Col4a3* (CKD), were monitored for a maximum of 12 weeks. **(A)** Gastrocnemius and tibialis anterior weights at endpoint (M-CTRL: n=15, M-CKD: n=14, F-CTRL: n=5, F-CKD: n=11). **(B)** Final grip strength (M-CTRL: n=7, M-CKD: n=7, F-CTRL: n=4, F-CKD: n=4). **(C)** Median CSA of muscle fibers and fiber distribution (gastrocnemius, M-CTRL: n=11, M-CKD: n=8, F-CTRL: n=4, F-CKD: n=6). Scale: 100 µm. **(D)** Muscle fibrosis (gastrocnemius, Sirius red; M-CTRL: n=10, M-CKD: n=9, F-CTRL: n=4, F-CKD: n=8). Scale: 100µm. **(E-F)** Gene expression of atrophy-related proteins in gastrocnemius (M-CTRL: n=8-9, M-CKD: n=9, F-CTRL: n=4-5, F-CKD: n=7-9): proteins involved in the ubiquitin proteasome pathway **(E)** and autophagy **(F)**. Data are mean ± s.e.m. Statistical analysis: unpaired (A-F, C *Median area*) or paired (C *Frequency*) two-way ANOVA with Sidak’s post-hoc tests. P values are displayed on the graphs. P < 0.05 was considered significant.

### Kidney failure is associated with adipose tissue loss and reduced lipid storage in Col4a3-deficient mice

Next, we assessed adipose tissue wasting as another determinant of cachexia, which is closely associated with survival in patients with CKD^2,25,26^. Overall loss of fat mass (**Figure 3E**) was confirmed by a strong reduction in the weights of gonadal and subcutaneous adipose tissue depots compared to CTRL (**Figure 5A**). Subcutaneous adipose tissue histology also confirmed the strong depletion in lipid stores, with a greater proportion of small adipocytes in animals with kidney failure (**Figure 5B-C**). At the molecular level, cachectic *Col4a3^-/-^* animals showed a reduction in total levels of *Acetyl-CoA Carboxylase* (ACC), a protein involved in lipid synthesis, and an increase in the lipolytic enzyme *Adipose Triglyceride Lipase* (ATGL) (**Figures 5D-E**). Despite sex-related differences (**Figures 3E, 5A**), we observed nevertheless a strong depletion in fat stores and similar molecular signatures in both males and females.

**Figure 5:**
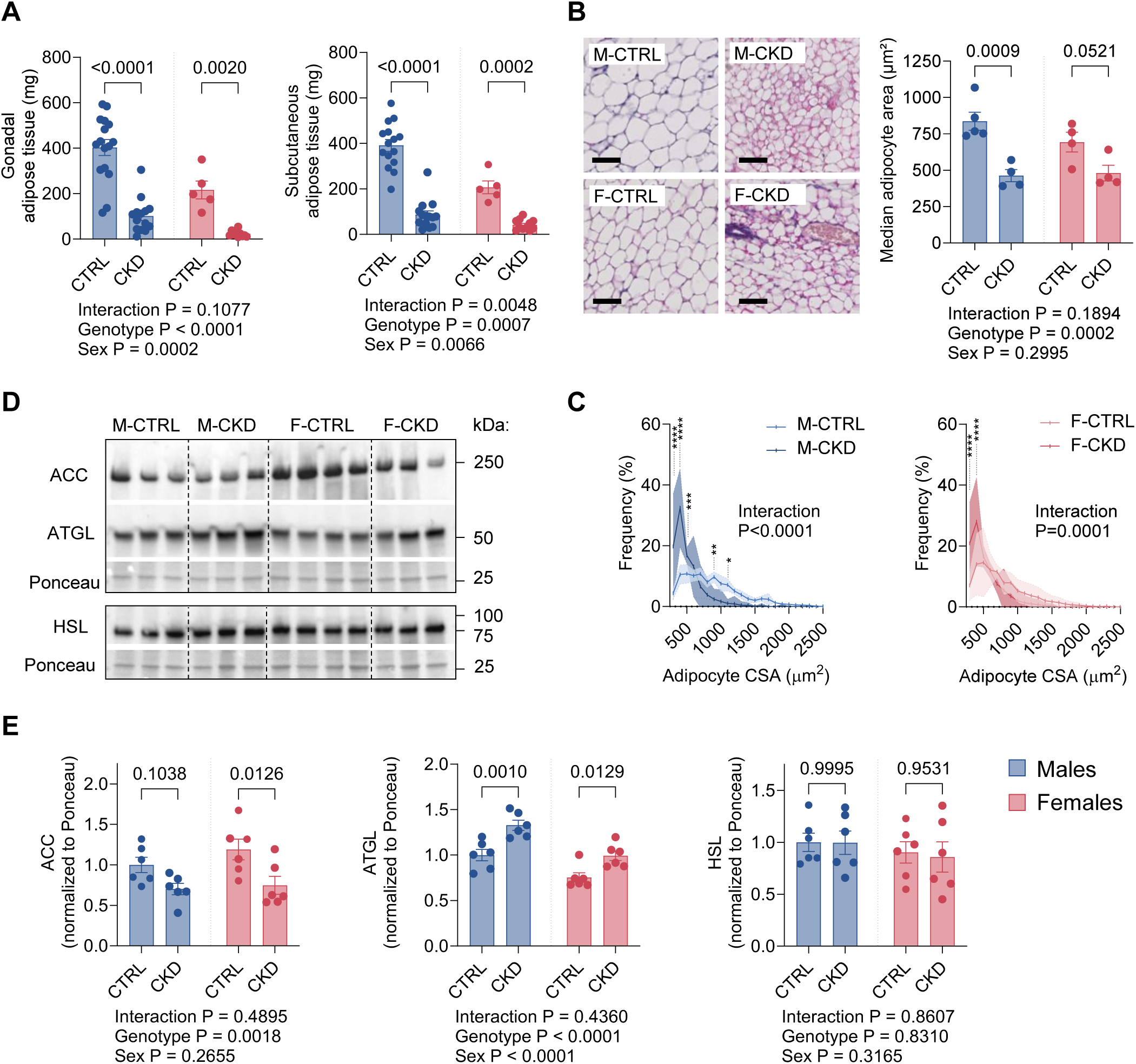
Kidney failure is associated with adipose tissue loss and reduced lipid storage in *Col4a3*-deficient mice. Male (M, blue) and female mice (F, red), heterozygous (CTRL) or knockout for *Col4a3* (CKD), were monitored for a maximum of 12 weeks. **(A)** Gonadal and subcutaneous adipose tissue weights at endpoint (M-CTRL: n=15-16, M-CKD: n=14-15, F-CTRL: n=5, F-CKD: n=10-11). **(B-C)** Median CSA of adipocytes in subcutaneous adipose tissue **(B)** and adipocytes’ distribution **(C)**. Scale: 50µm. (M-CTRL: n=5, M-CKD, F-CTRL, F-CKD: n=4). **(D-E)** Levels of proteins involved in lipid synthesis (ACC) and release (lipases: ATGL, HSL) (n=6 per group). Ponceau as loading control. **(D)** Representative images and **(E)** quantification. Data are mean ± s.e.m. Statistical analysis: unpaired (A, B *Median area*, E) or paired (B *Frequency*) two-way ANOVA with Sidak’s post-hoc tests. P values are displayed on the graphs. P < 0.05 was considered significant.

### Kidney failure is associated with systemic inflammation and the induction of pro-cachectic mediators in Col4a3-deficient mice

To further investigate the cachectic phenotype of *Col4a3^-/-^* animals, we assessed their pro-inflammatory status as a hallmark of cachexia. First, we measured gene expression of acute phase response proteins in liver and confirmed their strong induction in cachectic animals with kidney failure compared to CTRL (**Figure 6A**). We also confirmed elevated circulating levels of serum amyloid A (SAA, **Figure 6B**), an acute phase response protein with pro-atrophic potential on skeletal muscle^27^. In addition, we measured plasma levels of interleukin-6 (IL-6) and growth factor differentiation-15 (GDF-15), two well-known key pro-cachectic cytokines involved in muscle and adipose tissue wasting, liver dysfunction and anorexia in cancer-associated cachexia. Both were strongly induced in animals with kidney failure (**Figures 6C-D**), with comparable alteration in males and females.

**Figure 6:**
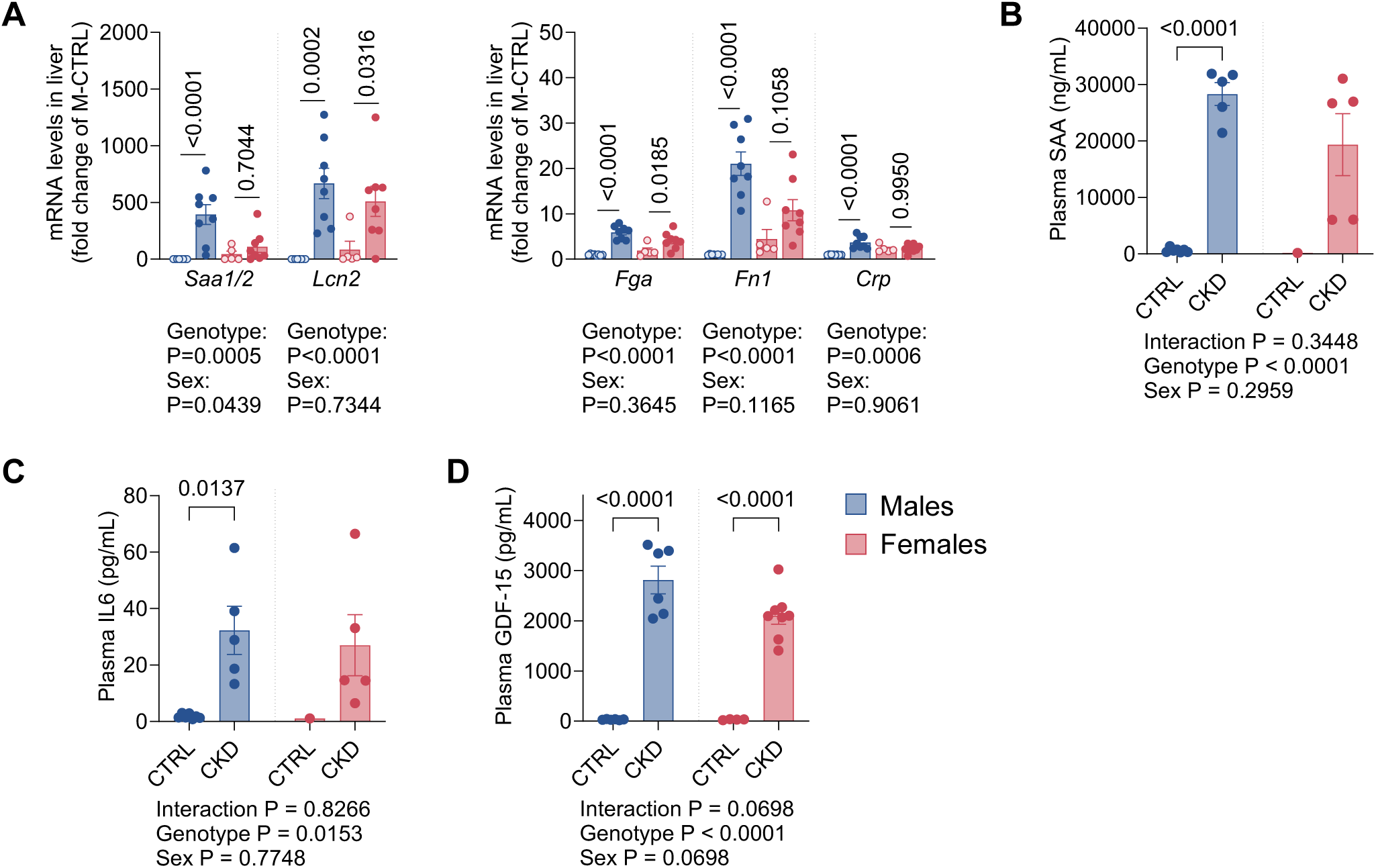
Kidney failure is associated with systemic inflammation and induction of pro-cachectic mediators in *Col4a3*-deficient mice. Male (M, blue) and female mice (F, red), heterozygous (CTRL) or knockout for *Col4a3* (CKD), were monitored for a maximum of 12 weeks. **(A)** Gene expression of acute phase response proteins in liver (M-CTRL, M-CKD, F-CKD: n=8, F-CTRL: n=5). **(B-D)** Plasma levels of pro-cachectic mediators: **(B)** Serum Amyloid A, **(C)** Interleukin-6 (M-CTRL: n=7, M-CKD: n=5, F-CTRL: n=1, F-CKD: n=5) and **(D)** Growth Differentiation Factor-15 (M-CTRL: n=6, M-CKD: n=6, F-CTRL: n=4, F-CKD: n=8). Data are mean ± s.e.m. Statistical analysis: unpaired two-way ANOVA with Sidak’s post-hoc tests. P values are displayed on the graphs. P < 0.05 was considered significant.

### Cachexia signature in muscle of Col4a3^-/-^ animals with kidney failure shares similarities with cancer cachexia

To obtain additional insight into the molecular mechanisms involved in muscle atrophy in renal cachexia, we performed transcriptomics of the tibialis anterior of cachectic *Col4a3^-/-^* animals. Muscle transcriptome underwent a complete remodelling during cachexia development as shown by principal component analysis and assessment of differentially expressed genes (**Figure 7A**). Pathway analysis revealed cachexia signalling (pathway #1, 8), autophagy (#3, 4) and pro-inflammatory signalling (#2) as top significant pathways with a positive z-score (≥1,5), supporting potential activation in renal cachexia (**Figure 7B**, left panel). It also highlighted potential effect of oxidative stress (#10) and pathways involved in the control cell cycle (#6, 7) and muscle mass (#5) as hallmarks of muscle atrophy in renal cachexia. On the other hand, pathways with negative z-scores (≤1,5) suggested an overall downregulation of extracellular matrix proteins involved in tissue organization (**Figure 7B**, right panel). To better understand the muscle phenotype of animals with renal cachexia in regards to other types of cachexia, we compared our dataset with recently published muscle transcriptome analysis from animals with cancer-associated cachexia^20^ (well-established mouse model of colon C26 carcinoma-induced cachexia). Strikingly, almost half of the genes differentially expressed in renal cachexia were regulated in a similar manner in cancer cachexia (**Figure 7C**). Accordingly, most of the top significant pathways with positive or negative z-score in renal cachexia were common with cancer cachexia (**Figure 7D**), suggesting a pro-atrophic effect on muscle. Looking for upstream regulators of muscle remodelling in renal cachexia, modulation of pro-inflammatory signalling (upstream regulators #2, 5, 7, 9, 10 on **Figure 7E**) was identified as the main driver of the molecular changes. Comparison with cancer cachexia revealed some disease-specific differences in the upstream regulators, with TGFβ being more important for cancer rather than for renal cachexia (**Figure 7F**). In accordance with existing literature on renal cachexia, a potential decrease in IGF1 signalling (z-score = -1) and the induction of *Aryl Hydrocarbon Receptor* (AHR) (z-score > 2), a receptor for uremic toxins, was also observed in muscle. Finally, FOXO proteins were also amongst the most significant upstream regulators, in accordance with pro-inflammatory signalling and the induction of atrogenes in muscle of *Col4a3^-/-^* animals (**Figures 7F, 4E**).

**Figure 7:**
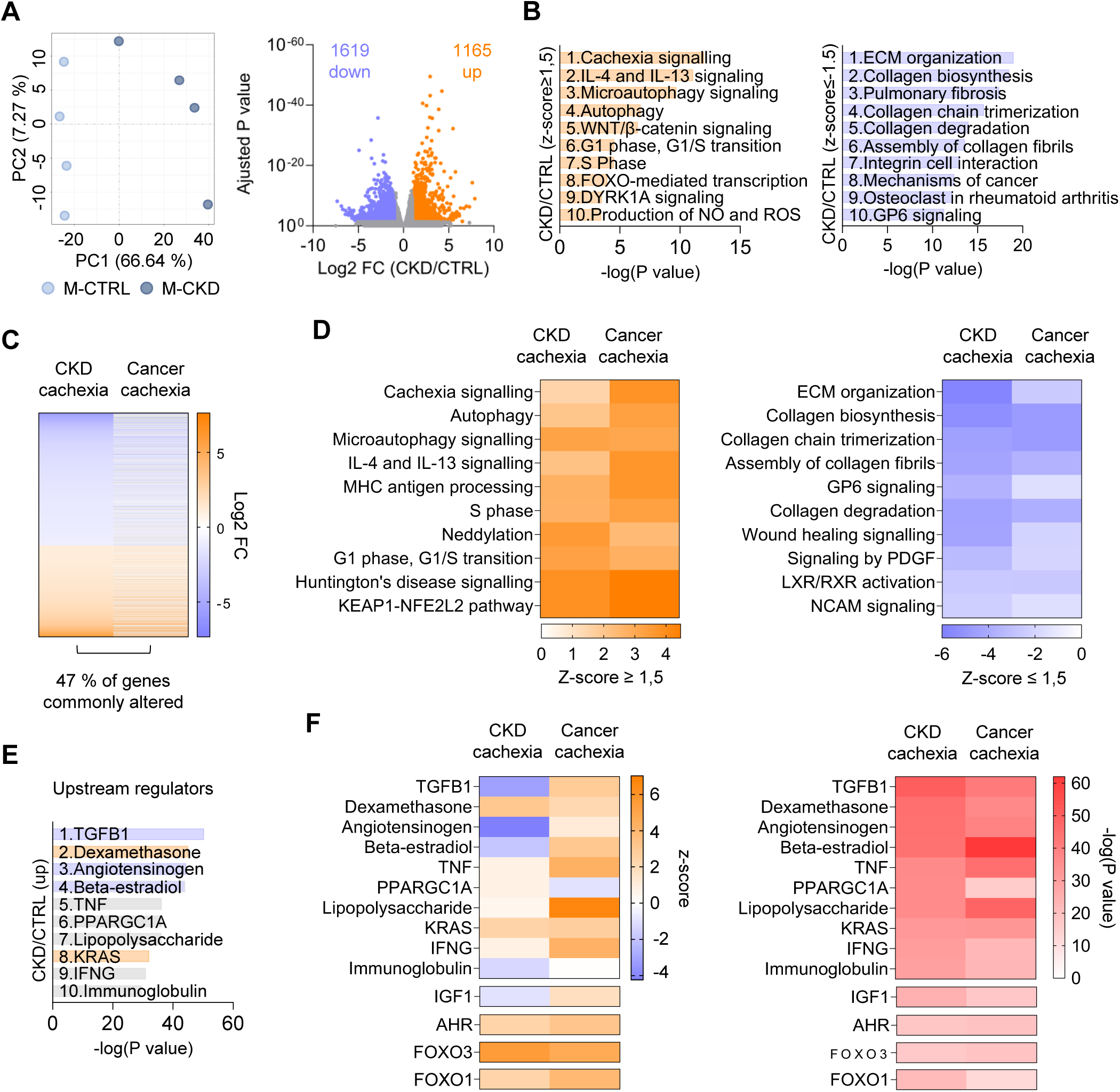
Cachexia signature in muscle of *Col4a3^-/-^* animals with kidney failure shares similarities with cancer cachexia. Transcriptomic analysis of tibialis anterior from control heterozygous (CTRL) and cachectic *Col4a3* knock-out male mice (CKD) (n=4 animals per group). **(A)** Principal component analysis of muscle transcriptome and volcano plot showing the n of genes altered in renal cachexia (FC > 1 or < -1 and ajusted P < 0.05). **(B)** Ingenuity Pathway Analysis (IPA, Qiagen). Top significant pathways with positive (orange) or negative (purple) z-scores (≥ 1,5 or ≤ -1,5). **(C-D)** Comparison of our dataset with muscle transcriptomics from animals with cancer cachexia (colon C26 carcinoma, gastrocnemius muscle) published in *Morigny et al, Nat Metab, 2026* ^20^. **(C)** n of genes commonly altered and **(D)** comparative pathway analysis highlighting top significant pathways with positive (orange) and negative (purple) z-scores (≥ 1,5 or ≤ -1,5). **(E-F)** Upstream regulator analysis (IPA, Qiagen). **(E)** Top significant upstream regulators in renal cachexia (orange: pathways predicted to be activated, purple: downregulated, and grey: no specified direction). **(F)** Comparison with cancer cachexia. In (F), both z-scores (left panel) and associated P values (right panel) are displayed. Data are mean ± s.e.m. Statistical analysis: please, see Methods’ section. Adjusted P < 0.05 was considered significant.

## Discussion

Cachexia is one of the strongest predictors of mortality in patients with kidney failure. Patients often experience frailty, reducing their quality of life and tolerance to therapies. Renal cachexia also predicts adverse cardiovascular and infectious outcomes and high hospitalisation rates. Beyond physical debility, renal cachexia also represents a major cause of emotional distress for patients and their families. Despite this strong impact, dedicated research is still scarce and there is currently neither standard of care nor approved therapies available.

In this study, we assessed the cachectic phenotype of the *Col4a3*^-/-^ mouse model of Alport syndrome, a well-established model of CKD^18^. Overall, our study highlights the potential of this model for the study of cachexia associated with kidney failure. Cachexia in *Col4a3*^-/-^ animals is defined by an ongoing loss of body weight, fat and lean mass together with the development of kidney failure. This model recapitulates the main features of cachexia, with the induction of proteolysis in muscle, lipolysis in adipose tissue, acute phase response in the liver and systemic inflammation. Transcriptomic analysis of skeletal muscle revealed a strong similarity with cancer cachexia and identified potential upstream regulators specific to CKD. In other mouse models of renal cachexia (nephrectomy, adenine), males are usually more susceptible to cachexia and muscle wasting^28^. Our analyses in the *Col4a3^-/-^* model did not reveal a strong sexual dimorphism on cachexia severity, although muscle wasting and fat loss seemed more pronounced in males.

An important limitation for the study of renal cachexia is the availability of CKD animal models with consistent cachectic phenotypes. Historically, the model of 5/6 nephrectomy, where the surgical reduction in total nephrons leads to a persistent decrease in GFR, has been the most referenced for the study of renal cachexia^1,4,10,11,13–17,28–31^. Alteration in body composition in this model englobes reduced weight gain compared to sham-operated controls rather than active weight loss, and is mostly explained by a reduction in fat mass gain. On the other hand, loss of lean mass has shown inconsistency between different studies, reporting either i) a loss of muscle mass, reduced grip strength and the induction of muscle atrogenes^14,28,31^, ii) unchanged muscle mass but reduced cross-sectional area associated with changes in fiber type, mitochondrial function, lipid infiltration and fibrosis^13,32^, iii) or no particular muscle phenotype^29^. The model of adenine-rich-diet-induced CKD is the second most used model for the study of renal cachexia. Adenine overload leads to tubular injury and to a decrease in GFR within 3 weeks associated with a body weight loss up to -30 % ^17,29,30^. With this setup, animals lose both adipose tissue and skeletal muscle mass associated with the induction of usual cachexia markers such as atrogenes. However, a main limitation in this model is the poor appetence of adenine-rich pellets, leading to severe anorexia^29^. Weight loss seems in consequence partly independent of CKD, and animals partially recover kidney function and body composition once diet is ended. Nevertheless, CKD in this model remains associated with long-term alteration in body composition, with persistent reduction in body weight and fat mass^29^. Therefore, those models are likely representative of pre-cachectic stages, where changes in body composition start to develop, but do not consistently meet the cachexia definition criteria which include an ongoing loss of body weight, lean and/or fat mass associated with functional impairment. Several approaches have been reported in order to circumvent this limitation and may represent good alternatives for the study of cachexia. For instance, feeding nephrectomised animals with protein-rich or phosphate-rich diets^33,34^, or using aged^35^ animals may promote stronger weight loss and/or muscle wasting. Furthermore, the combination of adenine with casein improves food appetence and reduces diet-induced anorexia, leading to the progressive development of kidney failure within 8 weeks and consistent cachectic phenotypes, which persist even two weeks after the end of the adenine-rich diet^28^.

The use of genetic models of CKD represents a good alternative for the study of cachexia without potential confounding effects resulting from surgery or diet. A model of polycystic kidney disease has been developed by extinguishing the expression of one of the main subunits of kinesin II (i.e. *Kif3a^-/-^*) in a kidney-specific manner. *Kif3a^-/-^* deficiency leads to the development of cysts, disrupting kidney structure and leading to kidney failure within 6 weeks^30,36^. In parallel, animals develop a weight loss of approximately 30 % associated with severe muscle wasting and weakness^30^. A disadvantage of this model however is the huge expansion of polycystic kidneys, which can represent up to 30 % of total body weight. One can therefore speculate that cachexia is in part due to the important compression of the gastrointestinal system, limiting food intake.

Our study highlights the potential of using the *Col4a3^-/-^*mouse model of Alport syndrome as a key model for the study of renal cachexia. It presents several advantages: i) the origin of CKD is kidney-specific without alteration in overall kidney size, ii) animals spontaneously develop kidney failure by the age of 8-12 weeks and recapitulate all main features of CKD^18,34^, iii) animals develop clear and highly reproducible cachectic phenotypes in response to kidney failure, characterized by an ongoing loss of body weight, lean and fat mass over 2-3 weeks once CKD develops, and iv) cachexia in *Col4a3^-/-^* mice is observed both in males and females, allowing for mechanistic insights into sex differences. One limitation may reside in off-target effects of whole-body knock-out of *Col4a3* in metabolic tissues. However, data from single-cell RNA-sequencing did not show any significant *Col4a3* expression in key metabolic cell types, including hepatocytes, myotubes or adipocytes. Alport patients do not suffer from growth development issues or specific metabolic alteration before reaching end-stage CKD^37^. In patients, *Col4a3* mutations can lead to sight and hearing disabilities, however whether this is the case in the *Col4a3^-/-^*model has not been studied.

Finally, one has to consider the importance of genetic backgrounds employed for the study of cachexia. In cancer, in animals injected with Lewis Lung Carcinoma (LLC), C57BL6/N mice are largely resistant to cachexia while C57BL6/J present fat and muscle wasting^35^. In nephrectomised or adenine-fed animals, C57BL6/J are often preferred, although the exact strain is not always mentioned. The FBV/N strain is more sensitive to CKD development, and could also be evaluated in regards to cachexia^38^. Similarly, while we report a strong cachectic phenotype of *Col4a3^-/-^*animals in the 129/SvJ strain, a similar deletion in the C57BL6/J background led to a delayed and milder impairment in kidney function and no particular cachectic phenotype^39^.

In conclusion, our study strongly encourages the use of 129/SvJ *Col4a3*^-/-^ animals for the study of cachexia associated with kidney failure. In this model, cachexia severity mirrors the severity observed in cancer, with high phenotypic and molecular similarities. This model will be particularly useful to identify joint altered pathways between cachexia of different pathological origins and molecular mechanisms specific to CKD-induced cachexia, and will help advancing our understanding of this complex, life-threatening, syndrome.

## Acknowledgements

We thank all the personnel from Inserm/UPS US006 CREFRE for their support with animal experiments. We thank I2MC core facilities (We-Met, Genotoul) for access to specific devices. We thank Mickaël Chabbert for his technical support with animals’ genotyping. We thank the PerMediK COST-Action CA21165 consortium for Personalized Medicine in Chronic Kidney Disease: improved outcome based on Big Data. LK, SB and JPS are members of EUTox.

## Funding

P.M. is funded by HORIZON-MSCA-2024-PF-01 (project #101202047 StopKidneyCachexia). This research was partially funded by a grant from the ANR PRCE programme (treatCKD, ANR-21-CE18-0031), the ANR JCJC programme (Calprotect, ANR-21-CE14-0013-01) and EP-Permed (EU-Calprotect, EPPERMED2024-099). M.R. is funded by the European Research Council (ERC) under the European Union’s Horizon 2020 research and innovation program (# 949017), an EFSD/Novo Nordisk Foundation Future Leaders Award, the DZD Next Young Talent Award, the German Research Foundation (DFG FOR5795-1_HyperMet), and the DZG Innovation Fund. L.K. is beneficiary of the European COST Action CA21165 and was funded by the French National Research Agency (ANR-25-CE14-0751-01). D.L. acknowledges the support of the European Research Council (ERC) under the European Union’s Horizon 2020 research and innovation programme (SPHERES, ERC Synergy Grant agreement No 856404).

## Ethics statement

All animal experiments were performed in compliance with the National Institute of Health’s Guide for the care and use of laboratory animals, 8^th^ edition, as well as the guidelines from the French Institute of Health for the care and use of laboratory animals. The project was approved by the local ethics committee for animal experiment (Inserm/UPS US006 CREFRE, CEEA122; Toulouse, France) and authorized by the Research French Ministry of Research and Higher Education (APAFIS #43673-2023053009321707).

## Conflict of interest

L.K. has received research grants from Fresenius Kabi, Nestlé, Lallemand, AstraZeneca, and Pfizer, as well as consulting fees, speaker honoraria, and travel support from AstraZeneca, Lilly, Baxter, Bayer, Novo Nordisk, and Fresenius Kabi.

---

**Figure S1:** Male (M, blue) and female mice (F, red), wild-type (WT) or heterozygous for *Col4a3* (HET), were monitored for a maximum of 12 weeks. Animals were age-matched (unless stated otherwise, M-WT: n=7, M-HET: n=16, F-WT: n=5, F-HET: n=5) **(A)** Age, **(B)** mGFR (M-HET: n=15, F-HET: n=4), and **(C)** kidneys’ weight at endpoint. **(D)** Initial (6-week-old) and **(E)** final body weight (at endpoint). Data from HET mice are the same as in Figures S3A, 2A, S3B, S2B and 3B. Data are mean ± s.e.m. Statistical analysis: unpaired two-way ANOVA with Sidak’s post-hoc tests. P values are displayed on the graphs. P < 0.05 was considered significant.

**Figure S2:** Male (M, blue) and female mice (F, red), heterozygous (CTRL) or knockout for *Col4a3* (CKD), were monitored for a maximum of 12 weeks (unless stated otherwise, M-CTRL: n=16, M-CKD: n=15, F-CTRL: n=5, F-CKD: n=11). **(A)** Initial mGFR (6-week-old, M-CTRL: n=9, M-CKD: n=6, F-CTRL: n=3, F-CKD: n=8). **(B)** Initial body weight. **(C)** Initial lean and fat masses (M-CTRL: n=7, M-CKD: n=7, F-CTRL: n=4, F-CKD: n=4). **(D)** Initial grip strength. Data are mean ± s.e.m. Statistical analysis: unpaired two-way ANOVA with Sidak’s post-hoc tests. P values are displayed on the graphs. P < 0.05 was considered significant.

**Figure S3:** Male (M, blue) and female mice (F, red), heterozygous (CTRL) or knockout for *Col4a3* (CKD), were monitored for a maximum of 12 weeks (unless stated otherwise, M-CTRL: n=16, M-CKD: n=15, F-CTRL: n=5, F-CKD: n=11). **(A)** Age at endpoint. **(B)** Kidneys’ weight at endpoint. Data are mean ± s.e.m. Statistical analysis: unpaired two-way ANOVA with Sidak’s post-hoc tests. P values are displayed on the graphs. P < 0.05 was considered significant.

**Figure S4:** Male (M, blue) and female (F, red) *Col4a3*-knock out mice were monitored for a maximum of 6-12 weeks. Correlation analysis between kidney function (mGFR) and body weight loss. Statistical analysis: simple regression analysis. P values are displayed on the graphs. P < 0.05 was considered significant.

**Figure S5:** Gene expression of *Col4a3* in **(A)** kidney, **(B)** liver, **(C)** skeletal muscle and **(D)** white adipose tissue. Plots were extracted from publicly-available databases: (A-C) the Mouse Cell Atlas (https://bis.zju.edu.cn/MCA/index.html) and (D) the Single Cell Portal (*A single atlas of human and mouse white adipose tissue* - https://singlecell.broadinstitute.org/single_cell). (A) Gene expression in kidney is shown at 10 days after birth and adult stages. *Col4a3* expression was undetectable at any stage in liver and muscle (only data at adult stage are displayed here).

## Notes

### Competing Interest Statement

L.K. has received research grants from Fresenius Kabi, Nestle, Lallemand, AstraZeneca, and Pfizer, as well as consulting fees, speaker honoraria, and travel support from AstraZeneca, Lilly, Baxter, Bayer, Novo Nordisk, and Fresenius Kabi.

