## Supplementary figures for "Identification of the *Col4a3*^-/-^ mouse as a model of chronic kidney disease-associated cachexia"

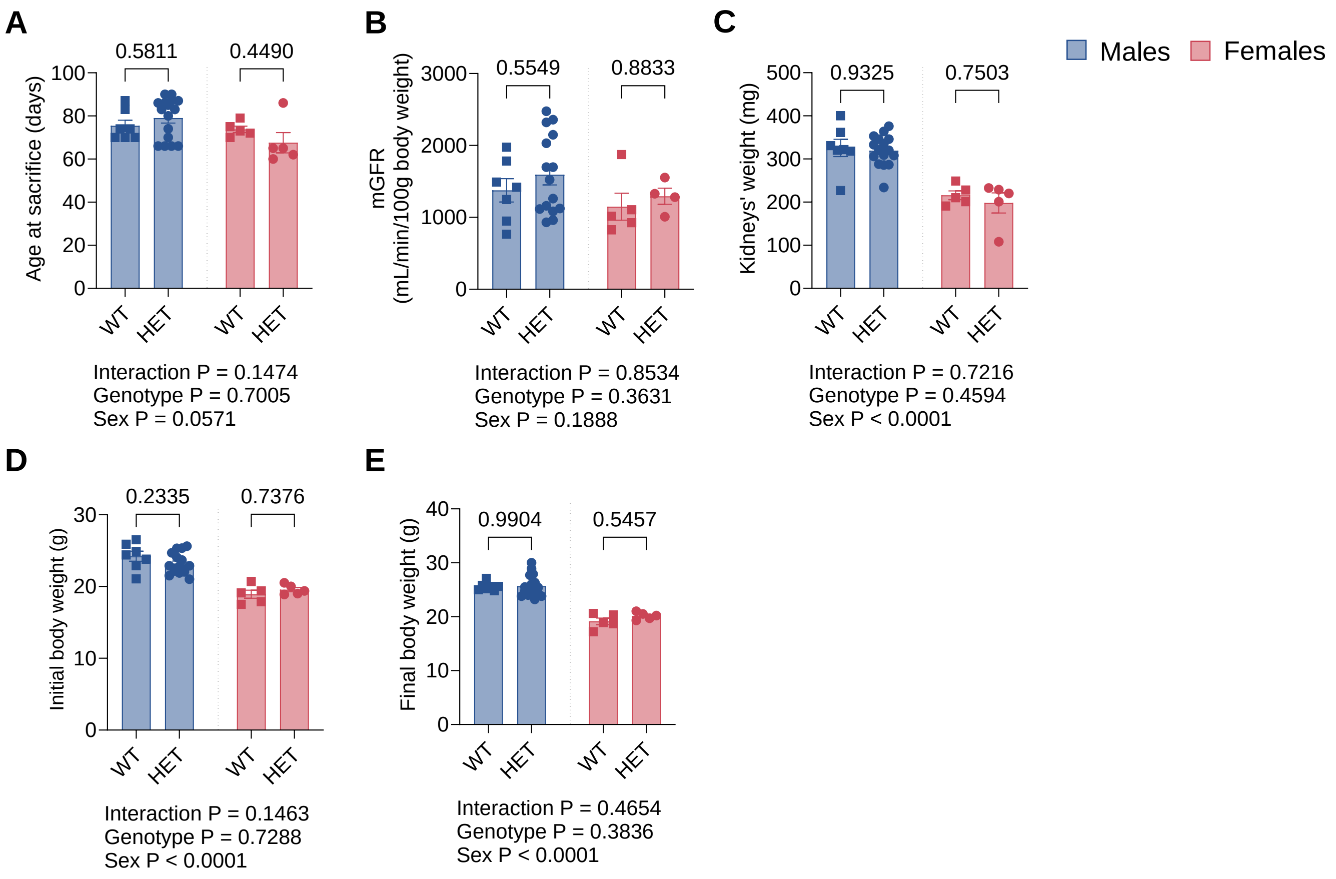


**Figure S1:** Male (M, blue) and female mice (F, red), wild-type (WT) or heterozygous for Col4a3 (HET), were monitored for a maximum of 12 weeks. Animals were age-matched (unless stated otherwise, M-WT: n=7, M-HET: n=16, F-WT: n=5, F-HET: n=5) **(A)** Age, **(B)** mGFR (M-HET: n=15, F-HET: n=4), and **(C)** kidneys' weight at endpoint. **(D)** Initial (6-week-old) and **(E)** final body weight (at endpoint). Data from HET mice are the same as in Figures S3A, 2A, S3B, S2B and 3B. Data are mean ± s.e.m. Statistical analysis: unpaired two-way ANOVA with Sidak's post-hoc tests. P values are displayed on the graphs. P < 0.05 was considered significant.

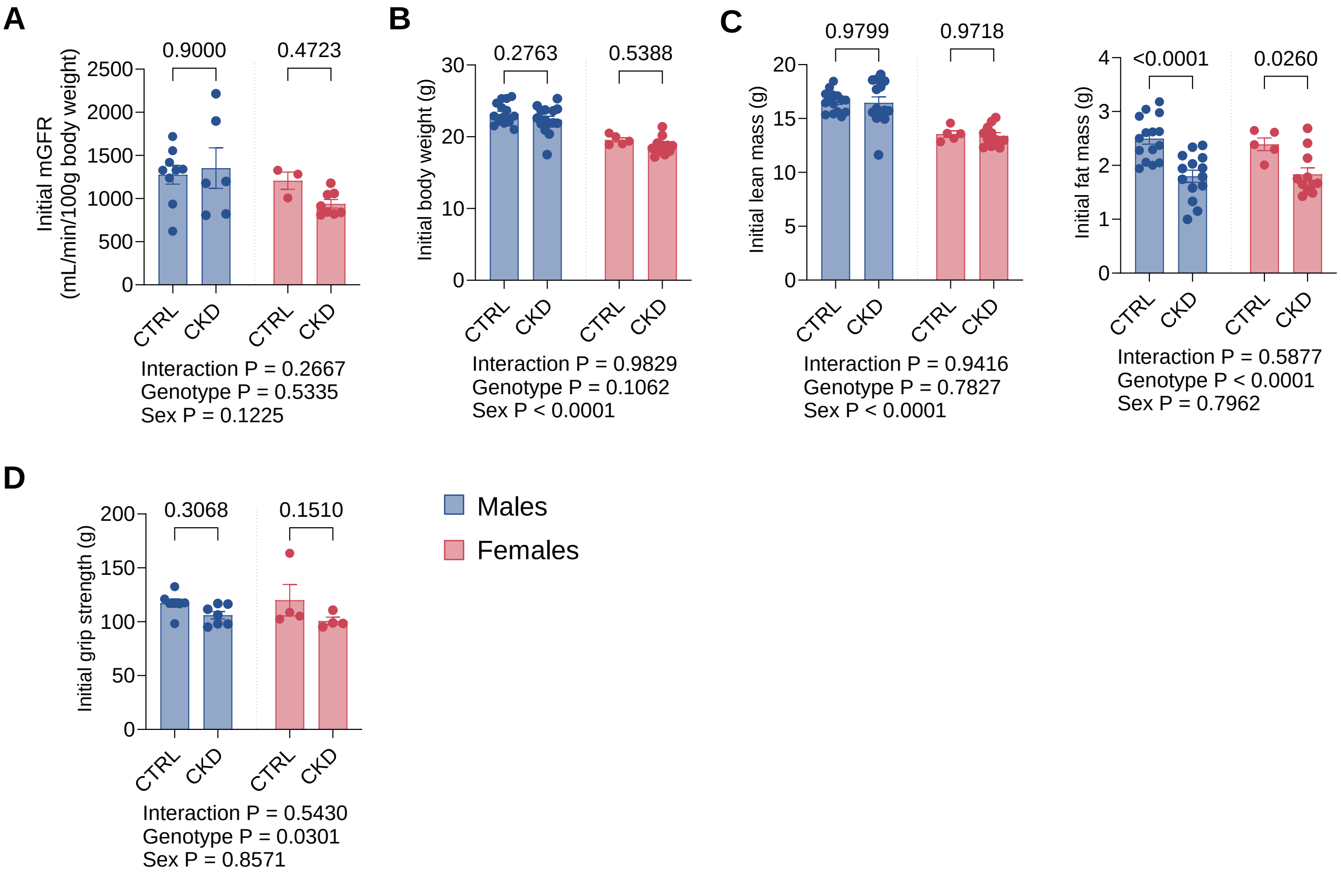


**Figure S2:** Male (M, blue) and female mice (F, red), heterozygous (CTRL) or knockout for Col4a3 (CKD), were monitored for a maximum of 12 weeks (unless stated otherwise, M-CTRL: n=16, M-CKD: n=15, F-CTRL: n=5, F-CKD: n=11). **(A)** Initial mGFR (6-week-old, M-CTRL: n=9, M-CKD: n=6, F-CTRL: n=3, F-CKD: n=8). **(B)** Initial body weight. **(C)** Initial lean and fat masses (M-CTRL: n=7, M-CKD: n=7, F-CTRL: n=4, F-CKD: n=4). **(D)** Initial grip strength. Data are mean ± s.e.m. Statistical analysis: unpaired two-way ANOVA with Sidak's post-hoc tests. P values are displayed on the graphs. $P < 0.05$ was considered significant.

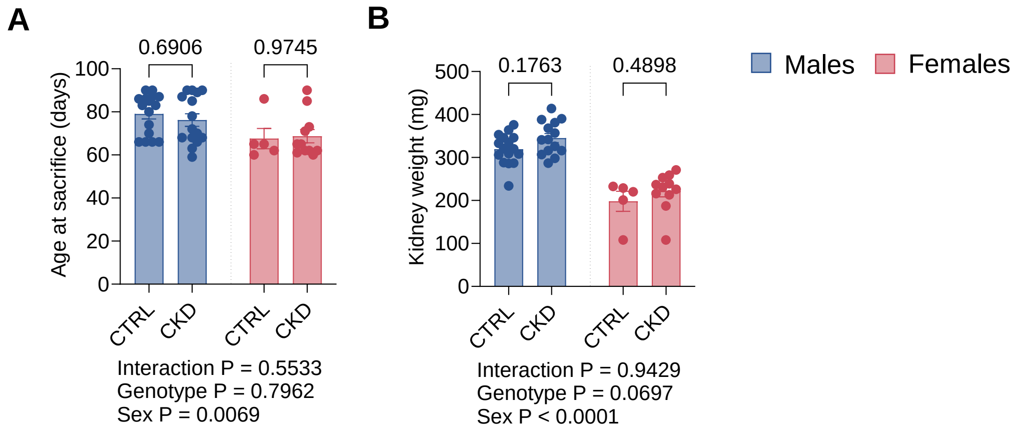


**Figure S3:** Male (M, blue) and female mice (F, red), heterozygous (CTRL) or knockout for Col4a3 (CKD), were monitored for a maximum of 12 weeks (unless stated otherwise, M-CTRL: n=16, M-CKD: n=15, F-CTRL: n=5, F-CKD: n=11). **(A)** Age at endpoint. **(B)** Kidneys' weight at endpoint. Data are mean ± s.e.m. Statistical analysis: unpaired two-way ANOVA with Sidak's post-hoc tests. P values are displayed on the graphs. $P < 0.05$ was considered significant.

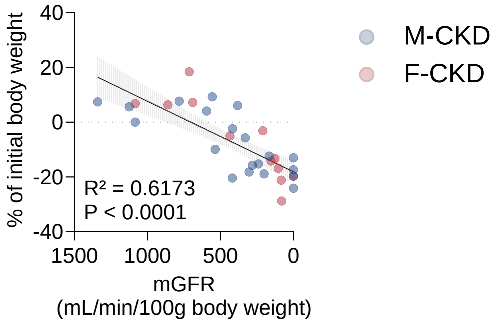


**Figure S4:** Male (M, blue) and female (F, red) Col4a3-knock out mice were monitored for a maximum of 6-12 weeks. Correlation analysis between kidney function (mGFR) and body weight loss. Statistical analysis: simple regression analysis. P values are displayed on the graphs. $P < 0.05$ was considered significant.

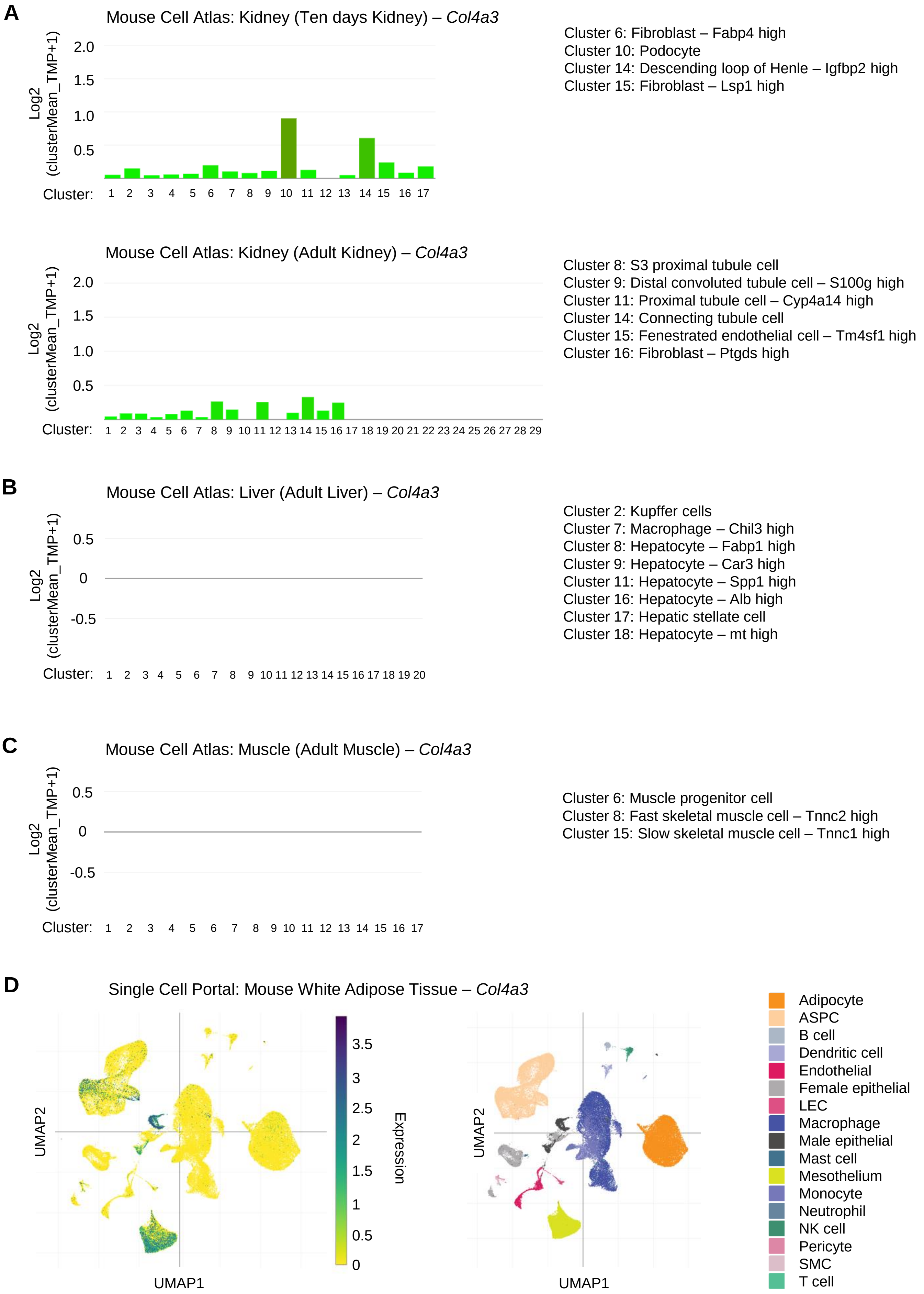


**Figure S5:** Gene expression of *Col4a3* in **(A)** kidney, **(B)** liver, **(C)** skeletal muscle and **(D)** white adipose tissue. Plots were extracted from publicly-available databases: **(A-C)** the Mouse Cell Atlas (https://bis.zju.edu.cn/MCA/index.html) and **(D)** the Single Cell Portal (*A single atlas of human and mouse white adipose tissue* - https://singlecell.broadinstitute.org/single_cell). **(A)** Gene expression in kidney is shown at 10 days after birth and adult stages. *Col4a3* expression was undetectable at any stage in liver and muscle (only data at adult stage are displayed here).
